# Ancient origin of Jingchuvirales derived glycoproteins integrated in arthropod genomes

**DOI:** 10.1101/2022.06.22.497255

**Authors:** Filipe Zimmer Dezordi, Gutembergmann Batista Coutinho, Yago José Mariz Dias, Gabriel Luz Wallau

**Author notes:** These authors contributed equally (Filipe Zimmer Dezordi, Gutembergmann Batista Coutinho, Yago José Mariz Dias).

## Abstract

Endogenous virus elements (EVEs) are viral-derived sequences integrated into their host genomes. EVEs of the Jingchuvirales order were detected in a wide range of insect genomes covering several distantly related families and Jingchuvirales-derived glycoproteins were recently associated by our group with the origin of a putative new retrovirus based on a glycoprotein captured by a mosquito retrotransposon. But, except for mosquitoes, there is a lack of a more detailed understanding of the endogenization mechanism, timing and frequency per viral lineages. Here we screened Jingchuvirales glycoprotein-derived EVEs (Jg-EVEs) in eukaryotic genomes. We found six distinct endogenization events of Jg-EVEs, that belong to two out of five known Jingchuvirales families (Chuviridae and Natareviridae). For seven arthropod families bearing Jg-EVEs there is no register of bona fide circulating chuvirus infection. Hence, our results show that Jingchuvirales viruses infected or still infect these host families, expanding their known host range. We estimated that two endogenization events occurred in the ancestors of the Myrmicinae-Ponerinae subfamilies (Pteromalidae - Hymenoptera order) around 155∼54.8 MyA and *Bombus* genus (Hemiptera order) around (36∼2 MyA). Although we found abundant evidence of LTR-Gypsy retrotransposons fragments associated with the glycoprotein in Hymenoptera and other insect orders, there is no evidence of potential functional glycoprotein capture. Our results show that the widespread distribution of Jingchuvirales glycoproteins in extant Arhtropods is a result of multiple ancient endogenization events and that these viruses’ fossils are being vertically inherited for millions of years through the Arthropods evolutionary tree.

**Graphical Abstract:** 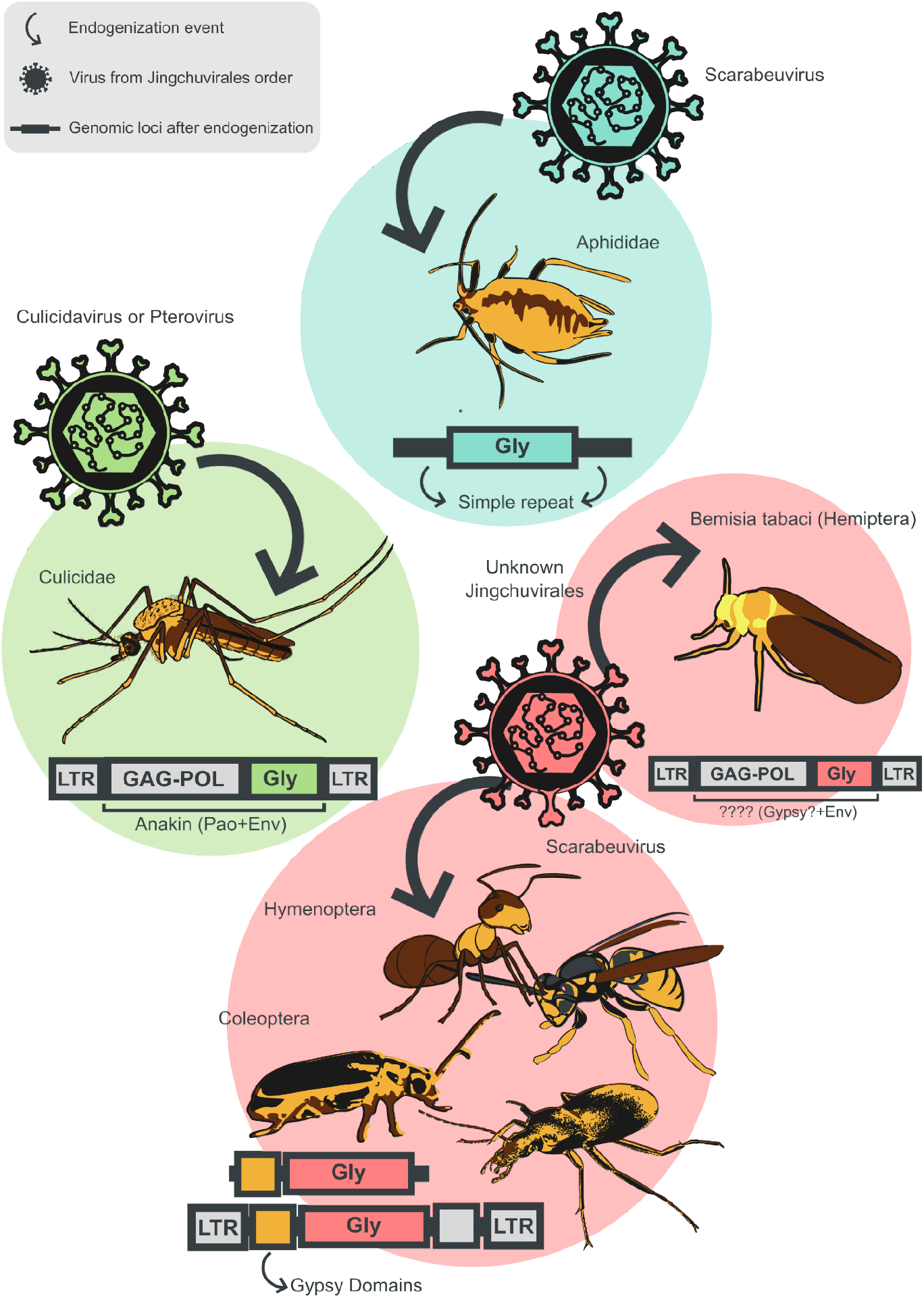

## Introduction

Viruses are the most abundant and diverse nucleic acid-based replicating units on Earth (Koonin & Krupovic, 2018). Viruses are parasitic replicating units that rely on infection and exploitation of cellular organism’s molecular machinery for their own replication. Because of this intimate and critical relationship with host cells, viruses and hosts undergo several interaction steps, even at the genome level (Coffin et al., 2021; Weiss, 2017). The retroviruses are known to integrate their genome into their host genome giving origin to Endogenous Retrovirus (ERVs). When integration takes place in germline cells ERVs are inherited by the next following host generations (Feschotte & Gilbert, 2012; Johnson, 2019). Interestingly, non-retroviral viruses also leave traces of past infection in their host genomes, and evidence of Non-retroviral Integrated RNA Virus Sequences (NIRVs), more broadly known as Endogenous Viral Elements (EVEs), can be found in a wide range of multicellular eukaryotic species (Katzourakis & Gifford, 2010). Increasing evidence shows that EVEs can be found in almost all host species investigated and that EVEs’ insertions may be neutral or impact the fitness of the organism (Armezzani et al., 2014; Ito et al., 2013; Ter Horst et al., 2019). Ultimately, the fixation of EVEs in a population or species depends on the endogenization frequency as well as their host fitness impact (Feschotte & Gilbert, 2012; Johnson, 2019).

The integration mechanism of non-retroviral sequences lacking retro transcriptases and integrases is still an open and intriguing question. It is notorious that the major part of EVEs identified in insects are derived from non-retroviruses (Gilbert & Belliardo, 2022). One of the most likely hypotheses is that integration is mediated by reverse transcriptase and integrases encoded by endogenous retrotransposons (Holmes, 2011; Katzourakis & Gifford, 2010). Retrotransposons are abundant and active in Insect genomes and may provide proteins *in trans* for viral cDNA synthesis and integration (Tassetto et al., 2019). Although integration may also occur through non-homologous recombination mediated by the double-strand break repair mechanism of the host (Katzourakis & Gifford, 2010). Moreover, there are clear discrepancies regarding EVEs viral families in Insects, that is, the most widespread and abundant EVEs derive from two viral families, Rhabdoviridae and Chuviridae known to infect Insects (Blair et al., 2020). It raises at least two interesting and related questions: Those viral families infect insects more frequently than other viral taxa increasing the chance of leaving abundant EVEs recorded in their host genomes? Do viral genomes from these families interact more frequently with endogenous retrotransposon proteins increasing their endogenization rate relative to other viral families? The availability of several Insect genomes and detailed characterization of EVEs may provide indirect evidence to explain these questions.

The Jingchuvirales order was first characterized in 2015 based on several complete genomes grouping into a large, well-supported and distinct monophyletic group of viruses found majoritarially from hosts of the orders Araneae, Neuroptera, Decapoda, Diptera and Ixodida (Li et al., 2015). These viruses are initially grouped into only one family (Chuviridae) with a negative-sense single-stranded RNA (ssRNA (-)) genome with distinct conformations such as unsegmented, segmented, linear or circular genomes (Li et al., 2015). Up to now, no viral isolation has been performed for viruses from that family and its description is restricted to viral genome sequences. In 2018, the International Committee on Taxonomy of Viruses (ICTV) created the Jingchuvirales order represented by the Chuviridae family only (Wolf Y et al., 2018) and more recently this family was split into 5 families and 19 genera based on RdRp similarity thresholds (Di Paola N et al., 2021). Interestingly, several homologous glycoprotein sequences of Chuviruses - the proteins that form the viral envelope – were found integrated into different host genomes, including mosquitoes (Dezordi et al., 2020; Li et al., 2015; Palatini et al., 2020; Russo et al., 2019; Whitfield et al., 2017), ticks (Li et al., 2015; Russo et al., 2019), flies (Li et al., 2015) and ants (Flynn & Moreau, 2019). Previous studies identified a higher number of endogenized glycoproteins of the Chuviridae family in different insect genomes when compared with endogenized nucleoproteins and polymerases (Russo et al., 2019; Whitfield et al., 2017). Our group recently showed that, in mosquitoes, such discrepancy occurred due to Chuviridae glycoproteins captured by endogenous retrotransposons followed by intragenomic replication and hence amplification of the glycoprotein sequences (Dezordi et al., 2020).

The new Jingchuvirales order and following family and genus level classification and the higher proportion of Chuviridae glycoproteins endogenized in insect genomes lead us to investigate a number of related questions in this study: Are there differences of Jg-EVE endogenization origin from the five Jingchuvirales families?; Are there specific associations of viral taxa (family/genus), host taxa and endogenous retrotransposons that explain the emergence and maintenance through the evolutionary time of Jg-EVEs?; What was the timing of endogenization events in the evolutionary history of Arhtropods? Here we performed an extensive literature review to catalog all complete genomes of Jingchuvirales order available, screened Jingchuvirales glycoprotein in eukaryotic genomes and reconstructed the phylogenetic history of complete genomes and endogenized glycoproteins. We detected that two out of five viral families of the order Jingchuvirales were involved in six ancient endogenization events and that all extant and widespread Jg-EVEs found in this study are derived from these ancient events.

## Materials and Methods

### Data collection

We performed a literature review using the database PubMed Central® (PMC). Initially, the identifying the published papers was carried out using the keywords “Chuvirus”, “Chuviridae” and “Jingchuvirales” and the boolean operator “OR” for the combination of these three terms. With the results from this search, we performed a screening based on reading the title and abstract. The papers that corresponded with the proposal were selected for a full reading. Finally, after the reading, the papers that, among the results, did not present relevant data to the research were excluded and those that remained were used in the study.

Only papers with the description of new chuvirus genomes, published up to May 2021 were selected. The exclusion criteria were papers describing only genomes from other viral families or with chuvirus genomes already available in previous papers; published before 2015, the year of publication of the first original chuvirus genomes, review articles, notes, and letters to the editor.

### Glycoprotein putative EVEs search

To identify putative Jingchuvirales glycoprotein-derived EVEs (Jg-EVEs) we used a BLASTp online approach. In this step, all Jingchuvirales glycoproteins identified in this study were used as queries against the non-redundant (nr) protein database updated in May 2021 excluding all viruses from the subjects. The results were clusterized to remove redundant hits using cd-hit with a sequence identity threshold of 100% (-c 1) and an alignment coverage threshold of 100% (-s 1) (Fu et al., 2012). The matching regions were reverse searched with correspondent genomes to select EVEs copies considering only matches with flanking regions of at least 10 kb. To analyze the Jg-EVEs boundaries, 10 kbp upstream and downstream flanking regions were extracted using the bedtools flank (Quinlan & Hall, 2010).

The flanking regions were used in three analyses to understand the genomic context of each Jg-EVEs. The repeat content was evaluated using the RepeatMasker (Chen, 2004) online tool (default parameters, DNA source: fruit fly) and the results were analyzed using an in-house R script (repeatmasker2heatmap.R) to evaluate the frequency of repeat classes into J-EVEs boundaries, this evaluation considered the total region length with some repeat or transposon signature by each Jg-EVE, and the region length of each repeat or transposon associated to the specific Jg-EVE. The same flanking regions were submitted to a domain signature analysis to identify putative hybrid elements originating from the capture of Jg-EVE by retrotransposons, where the ORFs were extracted using getorf (Gary Williams, 2000) (default parameters) and then analyzed with BATCH-CD-SEARCH (Marchler-Bauer & Bryant, 2004) (default parameters). Furthermore, we used cd-hit-est to identify clusters of Jg-EVEs plus flanking regions with a sequence identity threshold of 80% (-c 0.8), an alignment coverage threshold of 20% based on the longer sequence (-aL 0.2) in the most accurate mode of clusterization (-g 1) and then we used MAFFT (Katoh & Standley, 2013) with a global strategy to align the clusters to search for orthologous regions between arthropods genomes.

### Phylogenetic analyses

The RdRp protein was used to define the clades of the Jingchuvirales order and the glycoprotein was used to investigate the endogenization process across different eukaryotic groups. To reconstruct the RdRp phylogeny, RdRp from viruses belonging to Mononegavirales order according to ICTV (https://talk.ictvonline.org/) are recovered and clusterized (ictv_ncbi.py) from NCBI (https://www.ncbi.nlm.nih.gov/, last update at 2021 May) and are aligned separately by family. Each family alignment was automatically edited with CIAlign (Tumescheit et al., 2022), following the same strategy: an initial amino acid distance analysis followed by the automatic edition using the mean distance threshold for each family, the alignments were then concatenated and re-aligned. The RdRp of Jingchuvirales genomes bearing the three traditional proteins (glycoprotein, nucleoprotein, and RNA-dependent RNA-Polymerase) or with genome length equal to or greater than 9 kb are aligned with the Mononegavirales reference alignment. Phylogenetic analysis of glycoproteins was performed with alignments encompassing all reference chuvirus glycoproteins retrieved from the literature, glycoproteins recovered from a previous study (Dezordi et al., 2020) and glycoproteins retrieved through the aforementioned strategy.

Both nucleotide and amino acid alignments were performed with MAFFT, the substitution models are evaluated with ModelFinder (Kalyaanamoorthy et al., 2017), and the RdRp of Mono-chu sequences was reconstructed with MrBayes 3.2.7a (Ronquist et al., 2012) with two independent runs, stop value equals to 0.0049 and 25% of burnin. The glycoprotein of *bonafide* viruses and putative EVEs was used to reconstruct a phylogenetic tree using IQ-TREE2 (Minh et al., 2020). Branch support was assessed by the ultrafast bootstrap method (Hoang et al., 2018) with 1000 replicates. All trees were rooted using midpoint root and the annotation and visualization were performed using iTOL (Letunic & Bork, 2021).

## Results

Keywords searches on the literature databases resulted in 54 published studies (**Supplementary File 1**). After the full reading, only 21 met the inclusion criteria (**Table 1**). Since the first identification of chuviruses, 109 genomes associated with the Chuviridae family have been published of which 60 are complete genomes (carrying the tree hallmark proteins of the order - 73 nucleoproteins, 98 RNA-dependent RNA polymerase (RdRP), 79 glycoproteins **Supplementary File 3**). Of these, 49 were sequenced from samples of *Araneae, Blattodea, Decapoda, Diptera, Hemiptera, Ixodida, Neuroptera* orders which accounts for seven out of 26 known insect orders and *Perciformes* and *Squamata*, the remaining eleven from unspecified hosts (**Figure 1**).

**Table 1.** List of included studies that reported Chvirus genomes.

| ID*.Study | Year | New genomes | Countries | Host (Phylum) | Genome structure |
| --- | --- | --- | --- | --- | --- |
| 1. LI et al., 2015 | 2015 | 22 | China | Arthropoda | Circular: 17;<br>linear: 5 |
| 2. SHI et al., 2016 | 2016 | 18 | China | Arthropoda and<br>nematoda | Linear: 17<br>P-circular: 1 |
| 3. HANG et al., 2016 | 2016 | 1 | South Korea | Arthropoda | P-circular: 1 |
| 4. PINTO et al., 2017 | 2017 | 4 | Brazil | Arthropoda | Linear: 4; |
| 5. SOUZA et al., 2018 | 2018 | 8 | Brazil | Arthropoda | P-circular: 8 |
| 6. AIEWSAKUN;<br>SIMMONDS, 2018 | 2018 | 18 | China;<br>USA | Arthropoda | Circular: 15<br>Linear: 3 |
| 7. TOKARZ et al., 2018 | 2018 | 2 | USA | Arthropoda | Circular: 2 |
| 8. MEDD et al., 2018 | 2018 | 2 | Japan;<br>UK | Arthropoda | Linear: 2 |
| 9. SHI et al., 2018 | 2018 | 3 | China | Chordata | Linear: 3 |
| 10. HARVEY et al., 2019 | 2019 | 2 | Australia | Arthropoda | P-circular: 2 |
| 11. SAMEROFF et al., 2019 | 2019 | 2 | Trinidad and<br>Tobago | Arthropoda | Circular: 2 |
| 12. MAIA et al., 2019 | 2019 | 1 | Brazil | Arthropoda | Linear: 1 |
| 13. TEMMAM et al., 2019 | 2019 | 2 | Thailand | Arthropoda | Circular: 2 |
| 14. KÄFER et al., 2019 | 2019 | 33 | Germany;<br>Austria;<br>Venezuela;<br>France; China;<br>Japan;<br>Australia;<br>France; USA;<br>Unknown** | Arthropoda | Linear: 33 |
| 15. MISHRA et al., 2019 | 2019 | 1 | Saudi Arabia | Chordata | Linear: 1 |
| 16. THEKKE-VEETIL., 2020 | 2020 | 9 | USA | Arthropoda | Linear: 7<br>P-circular: 2 |
| 17. GONGARD et al., 2020 | 2020 | 2 | Guadeloupe;<br>Martinique | Arthropoda | Circular: 2 |
| 18. HAHN et al., 2020 | 2020 | 1 | USA | Platyhelminthes | Circular: 1 |
| 19. ARGENTA et al., 2020 | 2020 | 1 | Brazil | Chordata | Linear: 1 |
| 20. HAN et al., 2020 | 2020 | 1 | China | Arthropoda | Linear: 1 |
| 21. WU et al., 2021 | 2021 | 3 | USA;<br>Finland | Arthropoda | Linear: 3 |

\*Study ID used in this study to link with chuvirus genome present on **Supplementary File 2**, \*\*Genome with no geographic location available.

**Figure 1.**
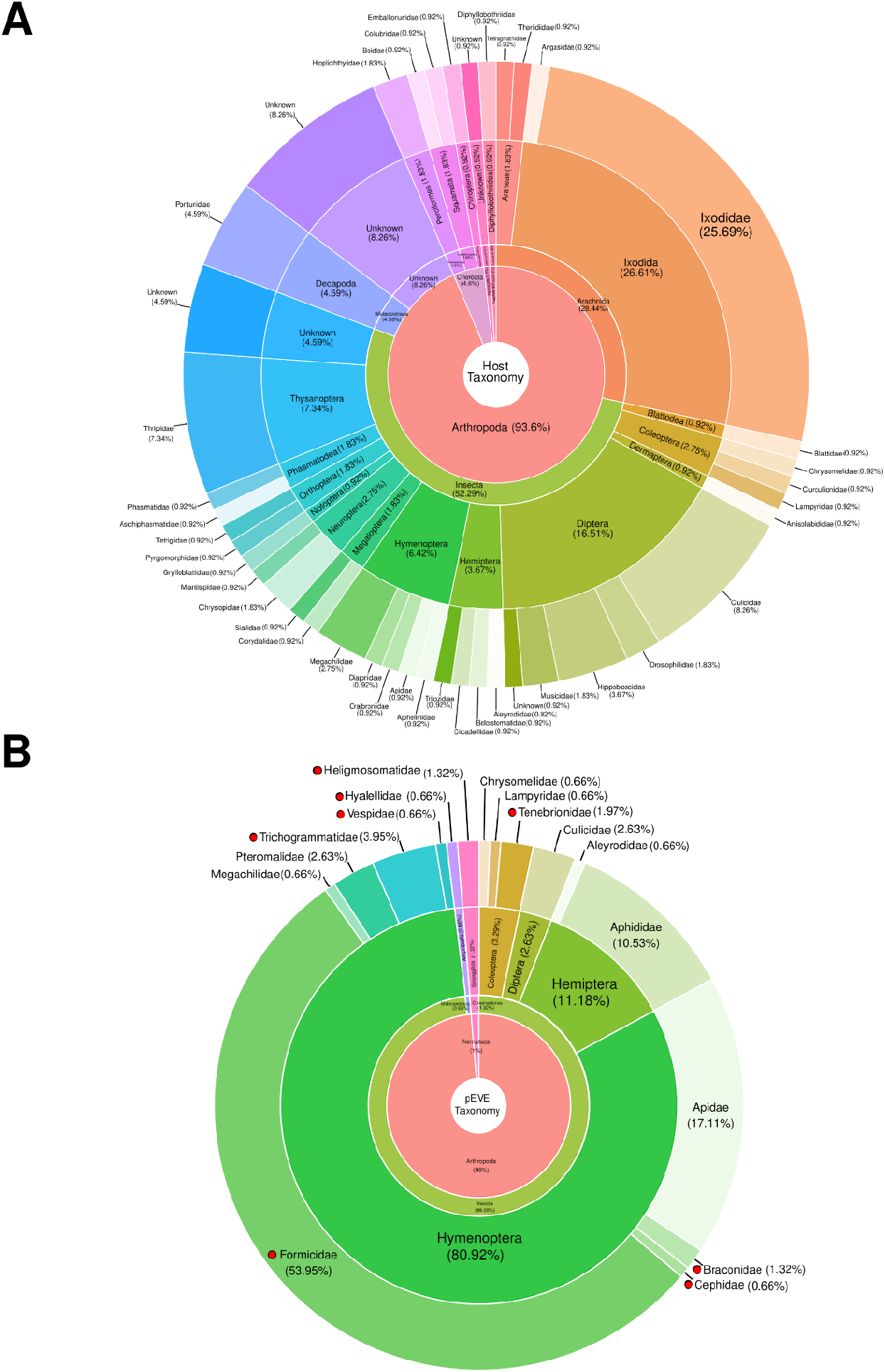
Host taxonomy of the Jingchuvirales order. **A**. Analyzed Jingchuvirales genomes (n = 109) sorted by different levels of host taxonomy. **B**. Jg-EVEs (n = 158), red circles represent host families where EVEs were found without a current register of bonafide Jingchuvirales infection in the literature. The percentage in each section of donut plots represents the total percentage of the section corresponding to the original entries of the innermost donut.

The RdRp phylogenetic tree confirmed the clustering of the new taxonomy of Jingchuvirales order proposed by ICTV (**Figure 2**), where the Chuviridae family comprises circular unsegmented genomes (Mivirus genus), circular bi-segmented genomes (Boscovirus), and diversity of linear or circular with 1 to 3 segments genomes (other 10 genera), the Aliusviridae, Myriaviridae, Crepuscuviridae and Natareviridae are represented by viruses with linear segmented genomes.

**Figure 2.**
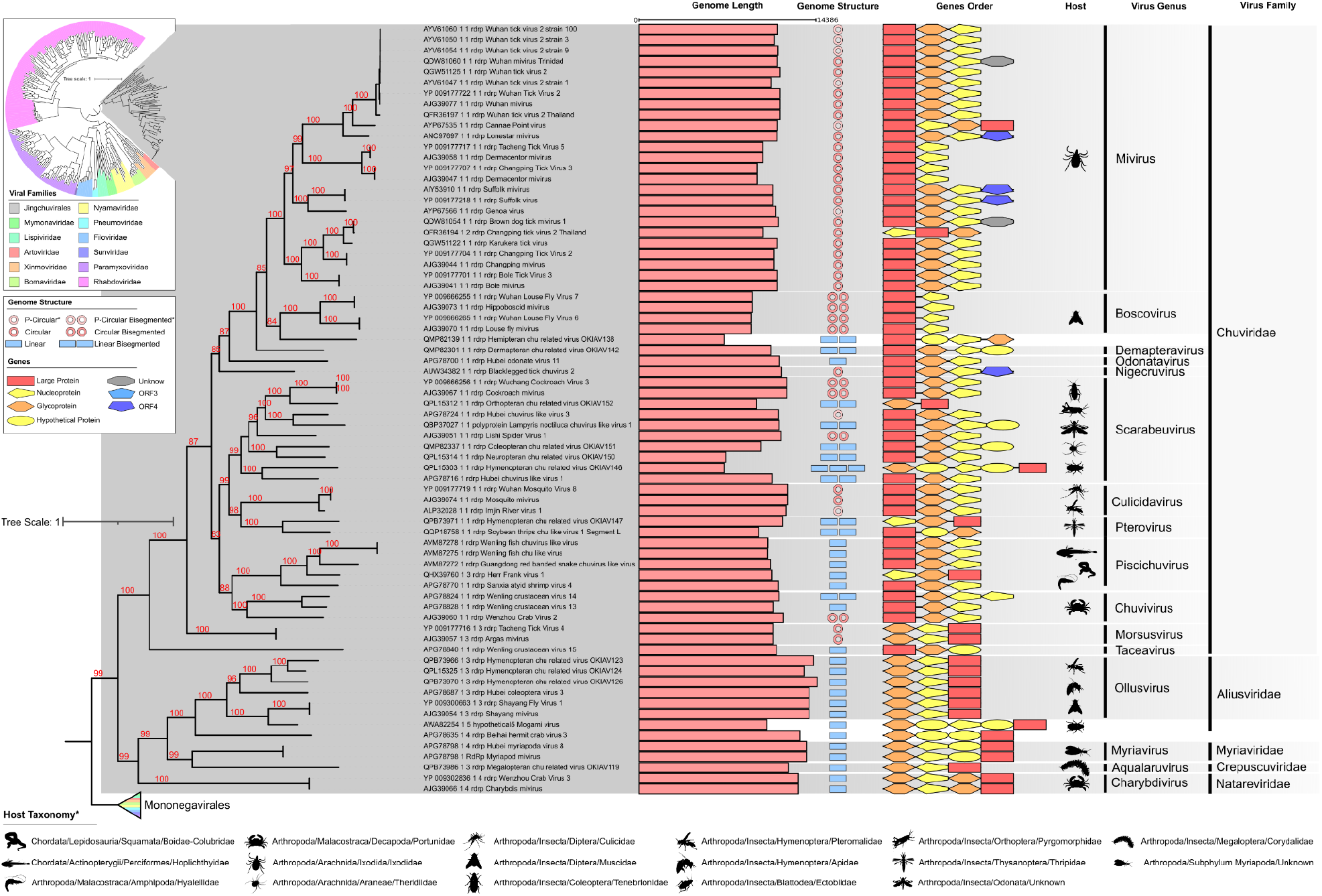
Bayesian phylogenetic tree of RdRp protein of Jingchuvirales and Mononegavirales orders. *Host Taxonomy updated at 2021 May according to NCBI information.

We found 158 EVEs (initial protein screening) representing 939 copies (screening against respective genome) found among 38 species (**Supplementary File 4**). The phylogeny of glycoproteins from Jingchuvirales and putative Jg-EVEs showed the existence of 6 distinct clades associated with endogenization events (**Figure 3**). From the six clades, one of them represents endogenization in Malacostraca with one Jg-EVE (**Figure 3B, Event-1**), one in Nematoda with two Jg-EVE (**Figure 3B, Event-3**), and four in Insecta (**Figure 3B, Event-2 4, 5 and 6**). We found Jg-EVEs of the different genera on the identified events. The Event-1 is related to the Chuvivirus and Piscichuvirus genus of the Chuviridae family, while Event-2, 5 and 6 are related to Pterovirus and other genera of the Chuviridae family, and the Event-3 to Charybdivirus genus (Natareviridae family) and the Event-4 to an unknown taxon of Jingchuvirales order.

**Figure 3.**
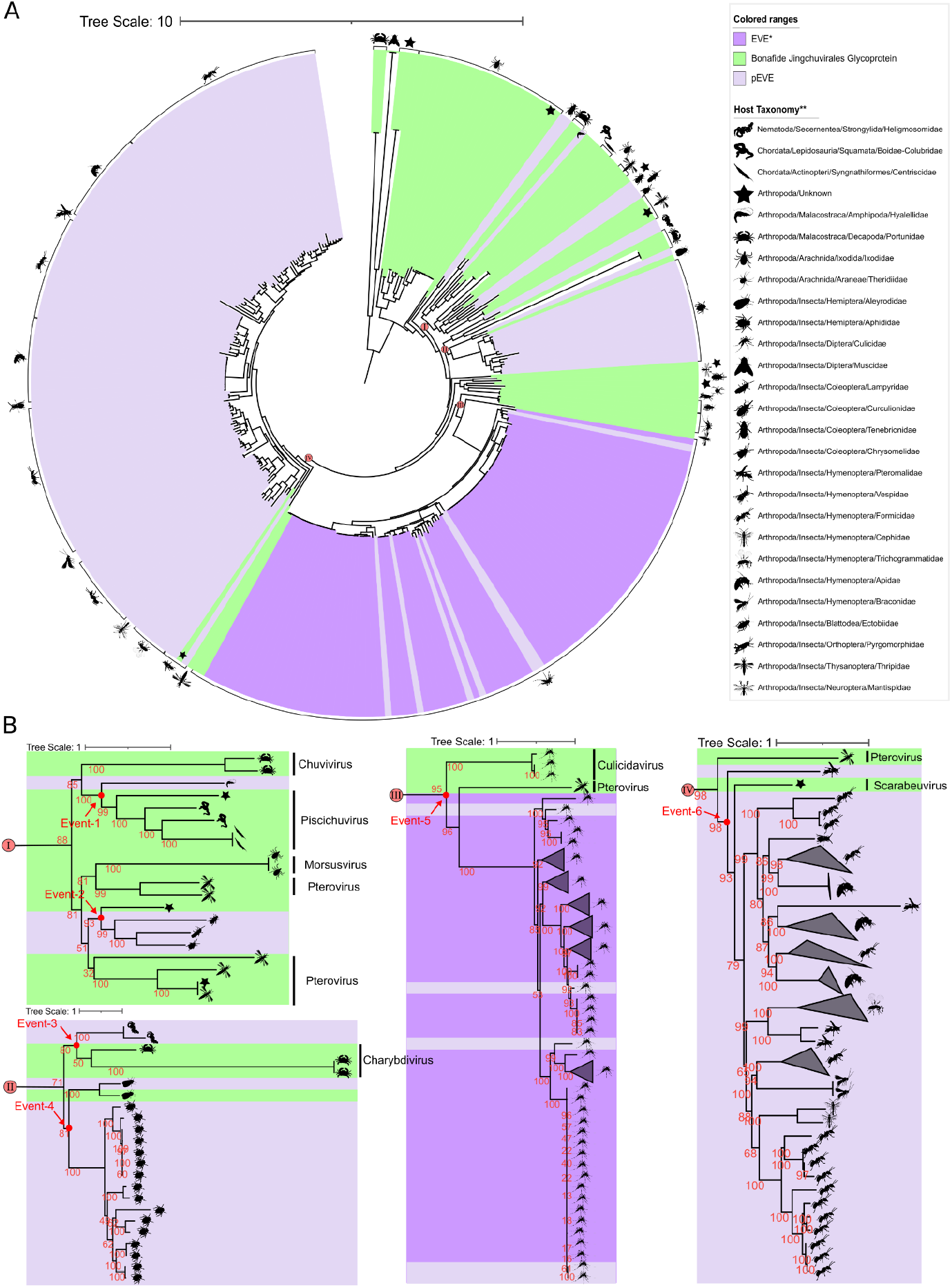
Maximum likelihood phylogenetic tree of the glycoprotein of Jingchuvirales order and putative representative Jg-EVEs. *EVEs recovered from Dezordi, et al 2020. **Host Taxonomy updated at 2021 May according to NCBI information.

**Figure 4.**
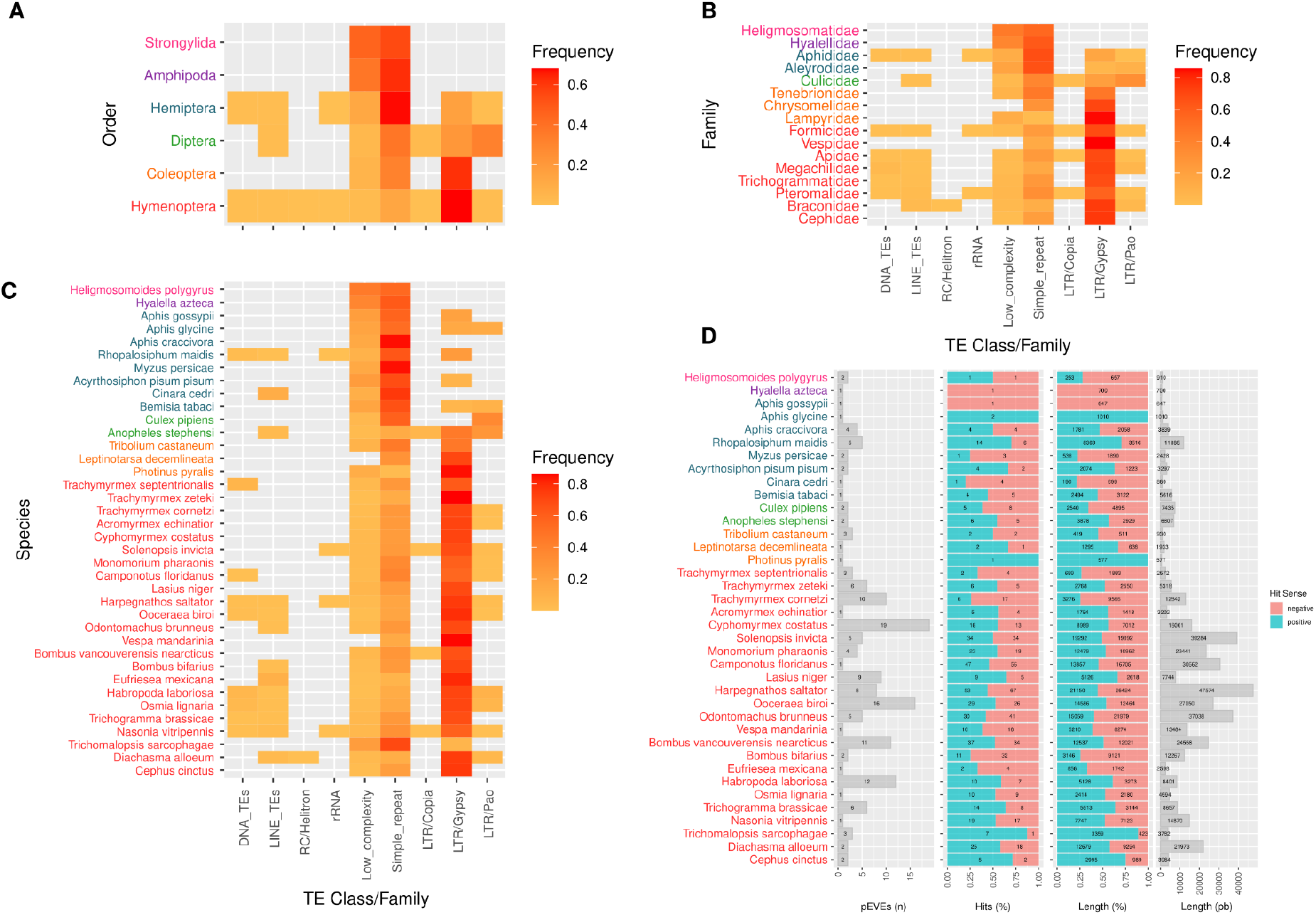
Jg-EVEs information. Heatmaps of frequency of TEs on EVEs boundaries at different taxonomy levels: (**A**) Order; (**B**) Family; (**C**) Species. (**D**) General information of representative EVEs hits against host genomes.

The Jg-EVEs of Malacostraca, Nematoda and some Insecta families are flanked by simple repeat and low complexity regions (**Figure 5**). On the other hand, several Jg-EVEs were flanked by LTR-Retrotranposon of Gypsy and BEL-Pao superfamilies in Hymenoptera, Culicidae and Coleoptera (**Figure 5**). The large majority of these associations occur between Jg-EVEs and fragmentary LTR retrotransposons copies (**Supplementary File 5**). However, we found two cases of Jg-EVEs in complete transposons boundaries, one in the species *Bemisia tabaci* (Unclassified element) and one Anakin (Dezordi, et al 2020) on *Anopheles stepehensi*.

Comparing a dataset of 939 Jg-EVEs copies and their flanking regions, we found 8 clusters with ortholog regions of different species (**Table 2**). The multiple alignments of each cluster showed conserved flanking regions between different species (**Supplementary File 6**), involving different species of the same genus (*Bombus* in clusters 263, 264, 265, 450, 451, 453 and 454) and from different genera of the same family (Pteromalidae in cluster 597). The endogenization event involving these Jg-EVEs occurred in the ancestral species of the *Bombus* genera around 36-2 MyA and in the ancestral of the Pteromalidae family around 155∼54.8 MyA (Kumar et al., 2017).

**Table 2.** Clusters of Jg-EVEs plus flanking regions.

| Cluster number | Identity (%) | Genus | Sequences | LCA |
| --- | --- | --- | --- | --- |
| 263 | 89.72-99.86 | *Bombus* | 4 | 36~2MyA |
| 264 | 83.64-94.28 | *Bombus* | 7 | 36~2MyA |
| 265 | 90.08-91.89 | *Bombus* | 3 | 36~2MyA |
| 450 | 81-57-86.88 | *Bombus* | 4 | 36~2MyA |
| 451 | 84.88-99.52 | *Bombus* | 4 | 36~2MyA |
| 453 | 82.85-87.06 | *Bombus* | 3 | 36~2MyA |
| 454 | 80.96-99.53 | *Bombus* | 10 | 36~2MyA |
| 597 | 80.24 | *Cyphomyrmex and Odontomachus* | 2 | 155~54.8MyA |

## Discussion

Viruses leave traces of past viral infection in their host genomes as EVEs (Patel et al., 2011). These elements have only recently received considerable attention and extensive characterization revealed that all known viral families can be found integrated into diverse host genomes (Blair et al., 2020; Feschotte & Gilbert, 2012; Katzourakis & Gifford, 2010). Insects are infected by a large diversity of viral families and cognate EVEs have been found in many genomes, but EVEs from two viral families are particularly enriched: Rhabdoviridae and Chuviridae (particularly glycoproteins) (Gilbert & Belliardo, 2022), raising questions about which host-viruses features may be generating EVEs endogenization disparities between viral families (Wallau, 2022). But the large majority of studies did not characterize EVEs in detail to be able to investigate such questions (Palatini et al., 2022). Based on previous findings regarding the capture and amplification of Chuvirus glycoprotein by a retrotransposon in mosquito genome we sought to investigate if the high content of Chuvirus glycoprotein EVEs also found in other Arthropod genomes could be derived from the same retrotransposon gene capture phenomenon and more broadly characterize the timing and number of events along with the Insects evolutionary history.

Our results showed four Jingchuvirales glycoprotein endogenization events in 38 eukaryote genomes investigated. Through the phylogenetic analysis, we were able to characterize four endogenization events that took place in the ancestors of several Insect taxa. Jg-EVEs widespread distribution in extant Insects may be a consequence of long-term by vertical transmission since those ancestral endogenizations. Ortholog copies of Jg-EVEs found between *Bombus* species (LCA 36∼2 MYA) and *Cyphomyrmex* (Myrmicinae subfamily) and *Odontomachus* (Ponerinae subfamily) ant genus (LCA 155∼54.8∼54.8 MyA) (Kumar et al., 2017). The orthologous regions provide further support for ancient integration events and long term vertical inheritance since the Eocene and lower Cretaceous Epochs, but studies focusing on high-quality genomes of specific host taxa should be performed to obtain more precise endogenization timing. All EVEs characterized were derived from two out of five currently recognized families of the order Jungchuvirales (Chuviridae and Netaviridae). Therefore, there is no specific association between EVE taxa and host taxa and the higher number of endogenization events derived from the Chuviridae family may be a result of its larger host range (**Figure 1**). Transposable elements and other repetitive sequences have been found in association with EVEs in several Insect species (Ter Horst et al., 2019; Whitfield et al., 2017). This evidence associated with experimental data shows that these repetitive endogenous host elements are mediating viral segment integration in the host genome (Tassetto et al., 2019) and that EVEs sequences and proteins may be co-opted as new genes of the host genome or captured by endogenous retrotransposons (Feschotte & Gilbert, 2012). Our analysis of Jg-EVEs showed that LTR retrotransposons of Gypsy, Copia, and Pao families are particularly enriched in their flanking regions. The first is highly prevalent in Hymenoptera and Coleoptera species while Copia and Pao are more clearly associated with Diptera species. However, despite such association, we found no clear evidence of Jungchuvirales glycoprotein capture by retrotransposons for Gypsy and Copia family other than the Pao retrotransposon capture of a Chuvirus-derived protein found in mosquitos by our group (Dezordi et al., 2020).

EVEs are equivalent to genetic fossils storing precious information about past viral infections and past or extant host range of specific viral families (Katzourakis & Gifford, 2010). Based on the phylogenetic relationships of the glycoprotein of *bona fide* viruses and Jg-EVEs clades, it is possible to infer that certain virus lineages may infect hosts previously not known to be infected by Jungchuvirals viruses. We found several EVEs in wasp (Vespidae) and ants (Formicidae) (**Figure 1B and Figure 3**), hosts that were not identified yet naturally infected by vírus from Jingchuviralres order (**Figure 2, Figure 1A**), but that have closely related taxa infected naturaly (**Figure 2**, Ollusvirus clade identified in Pteromalidae and Apidae, and Culicidavirus clade indenfied in Pteromalidae).

From the moment that the Chuviridae family was formally described (Li et al., 2015), the diversity of genomic structures was already known, with circular, linear, segmented and non-segmented genomes being observed, a particularity unusual in viruses. The authors proposed a phylogeny model in which it’s possible to observe an intermediate position of chuviruses between segmented and non-segmented genomes. However, it’s possible that such an organization follows an intrinsic evolutionary pattern within the Jingchuvirales order itself. Our RdRP phylogeny **(Figure 2)** shows different clade patterns, where we can notice clades exclusively with linear genomes and others with circular genomes. The new families Aliusviridae, Myriaviridae, Crepuscuviridae and Natareviridae proposed by ICTV are composed only of linear genomes arranged in specific clades of the Jingrchuvirales order (**Figure 2, Supplementary 7**).

In this study, we identified several Jg-EVEs across eukaryote genomes. These elements originated in the ancient past through six distinct integration events with the major part of events occuring on Insects. Despite the presence of TEs on EVEs boundaries, we found no evidence of glycoprotein capture by retrotransposons in other insect species except by the already characterized event in Culicidae. Therefore, new studies are warranted to better understand the deep relationships and long-term maintenance of Jingchuvirales glycoproteins EVEs in Insect genomes.

## Statements and Declarations

### Data availability

All supplementary tables are available on https://doi.org/10.6084/m9.figshare.20099549.v1, and a benchling notebook with all detailed steps is available on https://benchling.com/s/etr-bScSHJJqJ9f3GAc4xW5Y?m=slm-NSITX2bzgVQTXATDU8p6. All in-house scripts and phylogenetic files are publicly available on https://github.com/dezordi/jingchuvirales_dezordi_etal_2022.

## Author contributions

GLW and FZD conceived the study. GLW and FZD planned and supervised the work. FZD, GBC, YJMD carried out the analysis and wrote the manuscript. All authors agreed with the final version of the manuscript.

